# Seed-coat thickness explains contrasting germination responses to smoke and heat by *Leucadendron* species

**DOI:** 10.1101/2021.12.16.472977

**Authors:** Byron B. Lamont, Pablo Gómez Barreiro, Rosemary J. Newton

**Affiliations:** School of Molecular and Life Sciences (Ecology Section), Curtin University, PO Box U1987, Perth, WA 6845, Australia; Science Directorate, Royal Botanic Gardens Kew, Wakehurst Place, Ardingly, West Sussex, RH17 6TN, United Kingdom

**Keywords:** heat, *Leucadendron*, Mediterranean-type ecosystems, seed-coat thickness, seed permeability, serotiny, smoke, soil-stored seeds

## Abstract

Fire stimulates germination of most seeds in fire-prone vegetation. Fruits of *Leucadendron* (Proteaceae) are winged achenes or nutlets that correlate with their requirements for smoke and/or heat in promoting germination. We describe five possible smoke–heat dormancy-release/germination syndromes among plants, of which *Leucadendron* displays three (no response, smoke only, smoke and heat). As seed-coat thickness varies with seed-storage location (plant or soil) and morphology (winged or wingless), we tested its possible role in water uptake and germination. Species with winged seeds achieved 100% germination in the absence of smoke/heat, seed coats were an order of magnitude thinner, and their permeability greatly exceeded that of nutlets. As seed-coat thickness increased a) imbibitional water uptake declined at a decreasing rate, and b) the response to smoke, and to a lesser extent heat, increased linearly to reach levels of germination approaching those of winged seeds. For species responsive to smoke and heat, there was no additive effect when applied together, suggesting that they may have promoted the same physiological process. By what mechanisms a) the smoke response is greater the thicker the seed coat, and b) smoke chemicals could increase water permeability to explain the non-additive effect of smoke and heat, warrant further investigation.

**Highlight:** We show *Leucadendron* seeds are either plant-stored with thin, highly permeable seed-coats that germinate readily; or soil-stored and the thicker their seed-coat, the lower their permeability and greater their need for smoke/heat to promote germination.

## Introduction

Seeds in temperate, fire-prone ecosystems are usually stored in the soil but, in some regions, on the plant as well (Rundel *et al*., 2018). Three germination syndromes can be recognized in response to a summer-autumn fire event. 1: Soil-stored seeds that are impermeable to water rely on fire-heat to render them permeable before they can germinate (Tangney *et al*., 2020). 2: Soil-stored seeds that are permeable to water rely on smoke-associated chemicals to promote germination (Moreira *et al*., 2010). 3 Seeds stored on the plant (serotiny) are released from their fruits/cones in response to fire (branch death). These are then exposed to the soil environment and germinate as soon as substantial rain falls and daily temperatures drop below a threshold (Lamont *et al*., 2020). These three syndromes are usually considered discrete but there are increasing reports of their overlap. For example, despite insulation by their supporting structures, serotinous seeds are exposed to heat during a fire and a heat pre-treatment may sometimes promote germination (Midgley and Viviers, 1990; Hanley and Lamont, 2000). Some serotinous seeds may even respond to smoke chemicals that are released from soil particles following rain (Preston and Baldwin, 1999; Brown and Botha, 2004). Of particular interest are the (unexpected) reports of responses by species with soil-stored seeds to both smoke and heat (Kenny, 2000; Morris, 2000; Mackenzie *et al*., 2016). None of these studies sought to explain how these two fire-related factors could interact to break dormancy.

Newton *et al*. (2021) examined smoke and heat effects on the germination of 40 *Leucadendron* species. Fruits in this genus are single-seeded and thus can be treated as seeds; thus their pericarps are called seed coats. Untreated seeds of all but two of 31 serotinous species showed high levels of germination without any pre-treatment (mean of 96% at 20/10°C diurnal treatment in water agar, Table S1). Both serotinous species with poor control germination had nutlets rather than the winged achenes of the other serotinous species and either responded positively to aqueous smoke (*L. linifolium*) or germinated poorly (< 50%), irrespective of treatment (*L. album*). Smoke promoted germination in seven species that possessed nutlets (increasing from a mean of 24% among the controls to 92% germination when smoke-treated); of these, five were unaffected by 20 minutes of heat at 80°C and heat promoted germination in two. In other studies, germination of *L*. *daphnoides* and *L. tinctum* was quadrupled by scarification or removing the embryo from the seed (Brown and van Staden, 1973; Brown and Dix, 1985), consistent with physical barriers to germination. For the closely related *Leucospermum*, Brits and Manning (2019) showed that high temperatures resulted in desiccation and tearing of the endotesta that removed its impermeability to oxygen and enabled germination to occur.

Soil-stored nutlets can be expected to have thick seed coats to ensure resistance against decay agents, granivores and the digestive tract of animal dispersers (Calviño-Cancela *et al*., 2008; Hudaib, 2019; Dalling *et al*., 2020) and eventually fire heat, whereas serotinous seeds are protected and insulated by their supporting woody fruits or cones (Lamont *et al*., 2020). The latter can be expected to have thin seed coats since they germinate as soon as the soil is cool and moist. Thus, we wondered if seed-coat thickness might play a role in explaining the differences in germination requirements between these seed types. For example, water permeability decreases with increasing seed-coat thickness, independent of its hardness, in various legumes and grasses (Noodén *et al*., 1985; Frączek *et al*., 2005; Richard *et al*., 2018). Seeds that respond to fire-type heat usually have thick, dense, cutinized coats that are impermeable to fluids until heat opens up the dedicated ‘water gap’ in the seed coat (Moreira *et al*., 2010; Gama-Arachchige *et al*., 2013; Burrows *et al*., 2018). Among other species whose germination is promoted by heat (Hanley and Lamont, 2000; Kenny, 2000), no water gap is evident and general tearing of seed coat tissues seems to be involved (Brits and Manning, 2019).

Seeds that respond to smoke are weakly to moderately permeable (Moreira *et al*., 2010). This indicates a compromise in seed-coat thickness between protecting the embryo from deleterious agents in its environment and allowing smoke chemicals to reach the embryo. Thus, seed-coat thickness might provide a clue as to whether, and what sort of, a fire-related property is required to overcome embryo dormancy in *Leucadendron*. If the seed coat is thin, consistent with lack of soil storage and readiness to germinate as soon as dispersed, then the seeds will be highly permeable and there should be no fire response; if the coat is sufficiently thick to render it tardily permeable then the seeds will respond to heat; if the coat is moderately permeable then it will allow smoke chemicals to enter the seed and act catalytically (Flematti *et al*., 2004; Lamont *et al*., 2019) or increase its water permeability (Ghebrehiwot *et al*., 2008; Jain *et al*., 2008).

We therefore tested the following hypotheses:

1. The need for smoke and/or heat to promote germination of *Leucadendron* species is a function of seed-coat thickness;
2. Imbibitional water uptake is a negative function of seed-coat thickness and is inversely correlated with the promotory effect of smoke and/or heat on germination;
3. Serotinous seeds in winged fruits germinate readily in the absence if fire-related properties will have thin seed coats that are highly permeable to water, whereas nutlets that require smoke and/or heat to stimulate germination have relatively thick seed coats that are only weakly permeable.

## Materials and methods

The 40 species of *Leucadendron* examined are given in Table 1. The experimental design, involving controls, smoke, heat and (smoke + heat) pre-treatments, is described in Newton *et al*. (2021). *L. tinctum* showed low germination levels under all treatments and was not included in the analyses (but is considered in the Discussion). Five representative intact seeds were selected from the collections used for the germination experiments. Pericarps were bisected manually with a microtome blade. They were then positioned with the cut surface held horizontally under a Stemi dissecting stereoscope, model SV11, with a camera (AxioCam, Carl Zeiss, UK) attachment. Distance between outer and inner surfaces of the pericarp were measured in mm to 4 decimal places (AxioVision 4.8.1, Carl Zeiss). To examine the extent to which water permeability was a function of seed-coat thickness, 13 species were chosen to cover the three seed types and the full range of seed-coat thicknesses. Ten representative seeds per species were selected and their air-dry weights taken. Seeds were individually immersed in distilled water using a compartmentalized Petri dish, patted dry and weighed to 0.1 mg with a microbalance (UMT2, Mettler, Toledo) at 1, 3, 7, 24, 48 and 72-hour intervals, returning the seeds to the Petri dishes each time. Water content, expressed as a percentage, was determined as (wet weight minus dry weight)/dry weight and only the final result is reported here as only then had imbibition stabilized.

**Table 1.** 40 *Leucadendron* species whose seeds were subjected to simulated fire [viz., smoke (water) and/or heat (80°C for 20 min)] plus a dry, 40/20°C diurnal pre-treatment to simulate a warm postfire summer before incubation at optimal winter temperatures (moist, 20/10°C). Seeds plant-stored (P), or soil-stored (S). Fruit a nutlet (N) or flattened and winged (W). *L. nervosum* has small spindle-shaped achenes with long hairs that are unlike other nutlets listed here but is treated as W as it is wind-dispersed. Seed coat = arithmetic mean seed-coat thickness. Water content after imbibition for 72 h. SDs were 10-20% of the means and have not been included here. Germination data are posterior means. *Since this study was undertaken this species was found to be *L. galpinii*, although it was from a different collection, has slightly different propeties and is retained here.

| *Leucadendron* | Plant-<br>or soil-stored | Nutlet<br>or<br>winged | Seed<br>coat<br>(µm) | Imbi-<br>b<br>-ition<br>(%) | Germinatio<br>n<br>of controls<br>(%) |
| --- | --- | --- | --- | --- | --- |
| *album* | P | N | 200 | 37.0 | 36.2 |
| *argenteum* | P | N | 568 | 28.5 | 91.0 |
| *brunioides* | S | N | 316 | 38.5 | 3.6 |
| *chamalaea* | S | N | 228 | - | 81.1 |
| *comosum* | P | W | 24 | 57.3 | 99.8 |
| *coniferum* | P | W | 33 | - | 99.7 |
| *corymbosum* | S | N | 160 | - | 24.7 |
| *discolor* | P | W | 63 | - | 99.0 |
| *dregei* | P | N | 250 | - | 95.5 |
| *elimense* | S | N | 237 | 28.8 | 10.0 |
| *eucalyptifolium* | P | W | 39 | - | 100.0 |
| *flexuosum* | P | W | 48 | - | 100.0 |
| *foedum* | P | W | 44 | - | 100.0 |
| *galpinii* | P | N | 320 | 32.7 | 59.5 |
| *gandogerii* | P | W | 44 | - | 28.8 |
| *lanigerum* | P | W | 53 | - | 18.0 |
| *laureolum* | P | W | 40 | - | 100.0 |
| *laxum* | S | N | 234 | 34.1 | 28.8 |
| *linifolium* | P | N | 575 | 31.6 | 18.0 |
| *loranthifolium* | S | N | 154 | - | 24.9 |
| *meridianum* | P | W | 31 | - | 100.0 |
| *microcephalum* | P | W | 94 | - | 99.7 |
| *modestum\** | P | N | 236 | - | 63.5 |
| *muirii* | P | W | 24 | - | 99.4 |
| *nervosum* | P | N | 57 | 67.8 | 98.2 |
| *nobilis* | P | W | 27 | 55.5 | 99.8 |
| *procerum* | P | W | 70 | - | 100.0 |
| *rourkei* | P | W | 26 | 49.4 | 100.0 |
| *rubrum* | P | N | 51 | - | 99.8 |
| *salicifolium* | P | W | 58 | - | 99.8 |
| *salignum* | P | W | 47 | - | 100.0 |
| *sericeum* | S | N | 193 | - | 4.1 |
| *spissifolium* | P | W | 57 | - | 99.0 |
| *stelligerum* | P | W | 54 | - | 99.8 |
| *strobilinum* | P | W | 50 | - | 100.0 |
| *teretifolium* | P | W | 33 | - | 100.0 |
| *thymifolium* | S | N | 505 | 38.0 | 24.9 |
| *tinctum* | S | N | 955 | - | 48.1 |
| *uliginosum* | P | W | 140 | 40.9 | 88.1 |
| *xanthoconus* | P | W | 103 | - | 100.0 |

The statistical methods and allocation to smoke and/or heat responsiveness followed Newton *et al*. (2021). Germination data for the controls and treatment giving the highest mean result were extracted from Table S5 of Newton *et al*. (2021) and plotted against the seed-coat thicknesses obtained here. An increase in germination of 10% was selected as the minimum effect size of biological interest, with at least 95% confidence to separate non-trivial from trivial effects. Results with insufficient evidence to distinguish trivial from non-trivial effects were classified as ‘uncertain’ (Newton *et al*., 2021). The difference between germination levels of the controls and the three fire-related pre-treatments, and % water absorbed after 72 h, were plotted against mean seed-coat thicknesses. The lines with highest coefficients of determination (*R^2^*) among four curvilinear options [R (https://www.stats.bris.ac.uk/R) or Microsoft© Word for Mac 2011] were fitted to the data. Germination and seed-coat thickness data for species with winged, and non-winged plant or soil-stored seeds, or that did or not respond to fire-type properties were also grouped and compared by conventional ANOVA/Tukey’s statistics (http://vassarstats.net, R. Lowry©).

## Results

Thirteen species, all plant-stored with winged seeds and seed-coat thickness of 45 ± 19 μm (mean ± SD), germinated at ~100% among the controls. For the remaining 26 species (omitting *L. tinctum*), data for the controls fitted the linear equation: Y = −0.13X + 91.01 where Y = % germination and X = seed-coat thickness in μm (*R* = 0.543, *P* = 0.0041). Thus, estimated Y = 90.4% when X = 50 μm, and Y = 13.0% when X = 600 μm (the limit of seed-coat thicknesses recorded here). For the treatment yielding greatest response to smoke and/or heat, Y = 95.8% (*R* = 0.123, *P* = 0.5541), independent of seed-coat thickness, i.e., the treatment brought mean germination almost to the level of the 13 species not requiring any treatment to yield 100% germination.

Overall, the difference between germination levels of the controls and those treated with heat (Δ%) increased slightly with increase in seed-coat thickness, with zero difference at 23 μm and 10.7% at 600 μm (Fig. 1a). Addition of smoke increased the mean difference to 35.5% at 600 μm (Fig. 1b). Addition of smoke plus heat gave no further increase with a mean difference of 36.5% at 600 μm (Fig. 1c).

**Figure 1.**
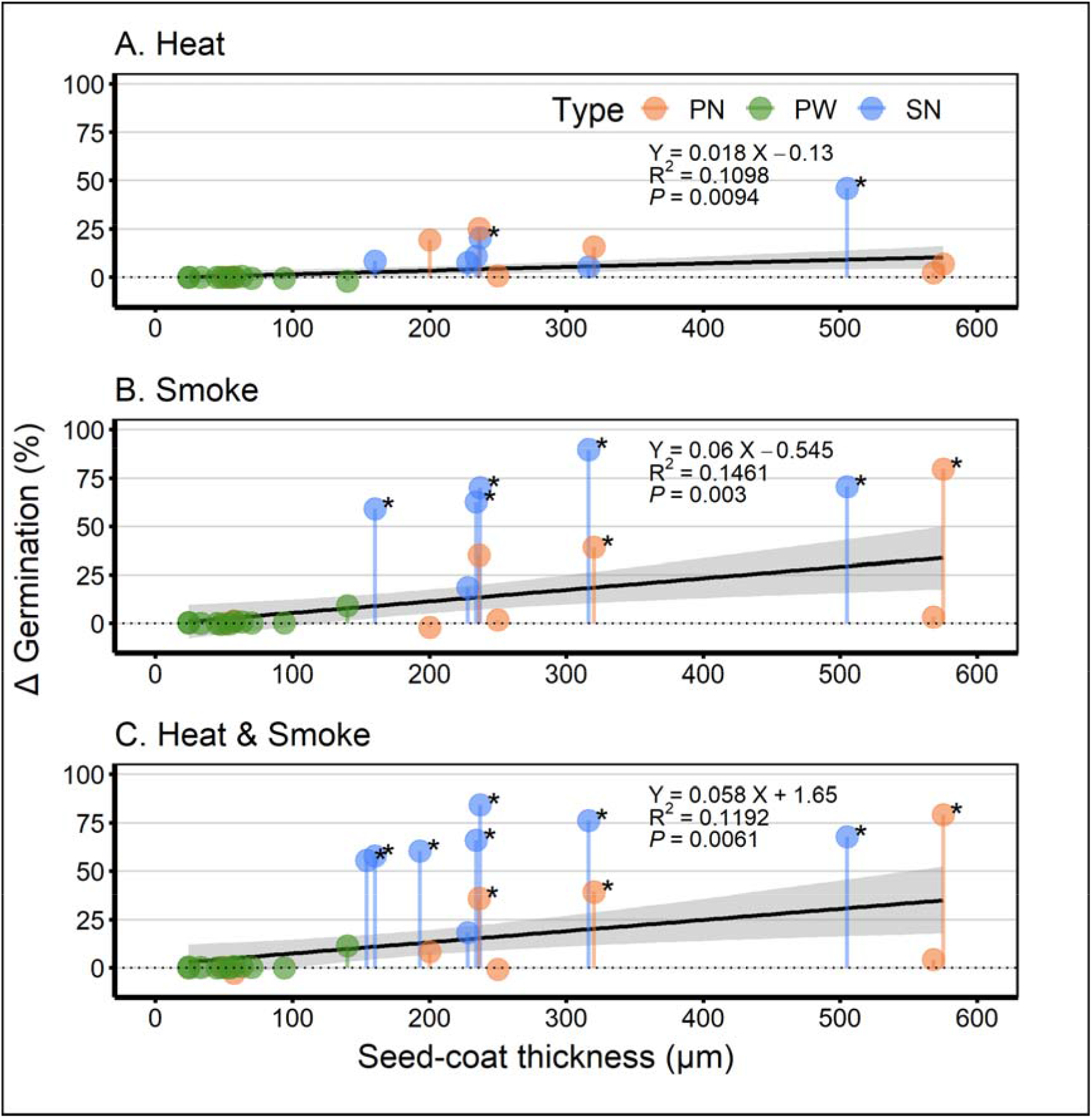
Relationship between seed-coat thickness and germination increase (Δ = difference between average treatment and average control germination) for all species whose controls were <100% germination. The 13 species with 100% germination among the controls are represented here by one value to minimize bias in the data. A. Following heat pre-treatment (80°C for 20 min), B. Following smoke water pre-treatment, and C. Following both heat and smoke. Seed types are Plant-stored Nutlets (PN, orange), Plant-stored Winged achenes (PW, green) and Soil-stored Nutlets (SN, blue). Biologically significant results (according to Newton *et al*., 2021) have an asterisk above/next to them. Linear regression lines are also given with their equations and probabilities, together with 95% confidence interval in grey. Germination averages were calculated from posterior mean germination values extracted from Table S5 of Newton *et al*. (2021)

The 13 species selected for study of imbibition showed similar trends to the total species in the study: these best obeyed a negative logarithmic function, with the controls ranging from ~100% germination at ~25 μm to ~ 10% at ~600 μm. As an index of permeability, seed water content after soaking for 72 h best declined in a power-function manner with increasing seed-coat thickness (Fig. 2). Estimated mean water content fell from ~60% at ~25 μm seed-coat thickness to ~42% at 140 μm (the range of thickness values for winged achenes), and ~50% at ~50 μm to ~30% at 600 μm (the range for nutlets).

**Figure 2.**
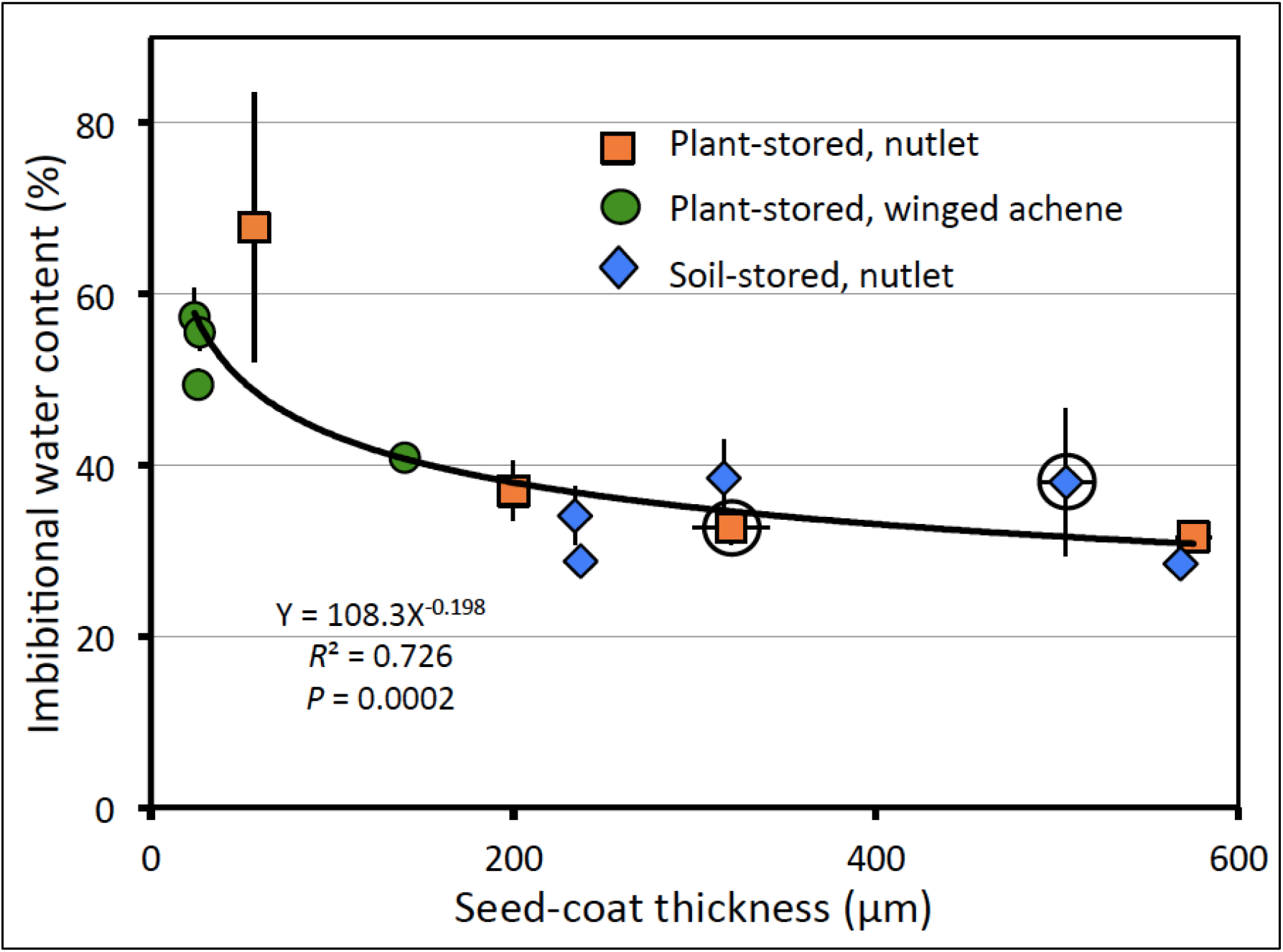
Mean ± SE imbibitional water contents after 72 h of soaking for the 13 species selected to represent the range of mean ± SE seed-coat thicknesses. The two species that responded to both smoke and heat are ringed. The best-fit curve to the data is given as well as its formula and probability.

Placing the data into the three location-morphology categories summarizes the previous results (Fig. 3). Germination responses to smoke (and heat) increased from winged to non-winged plant-stored seeds to soil-stored seeds. However, seed-coat thicknesses of the nutlets, whether plant- or soil-stored, were on average about six times thicker than the winged achenes (Fig. 3B). The only plant-stored seeds not to require a fire-related property for high germination levels were winged (74%); the only plant-stored seeds benefitting from smoke and/or heat were nutlets (10%) and their seed coats were > 7 times thicker on average, although with a large error term (Fig. 3C). All but 12% (one species uncertain) of the soil-stored nutlets responded to fire-related properties, particularly smoke.

**Figure 3.**
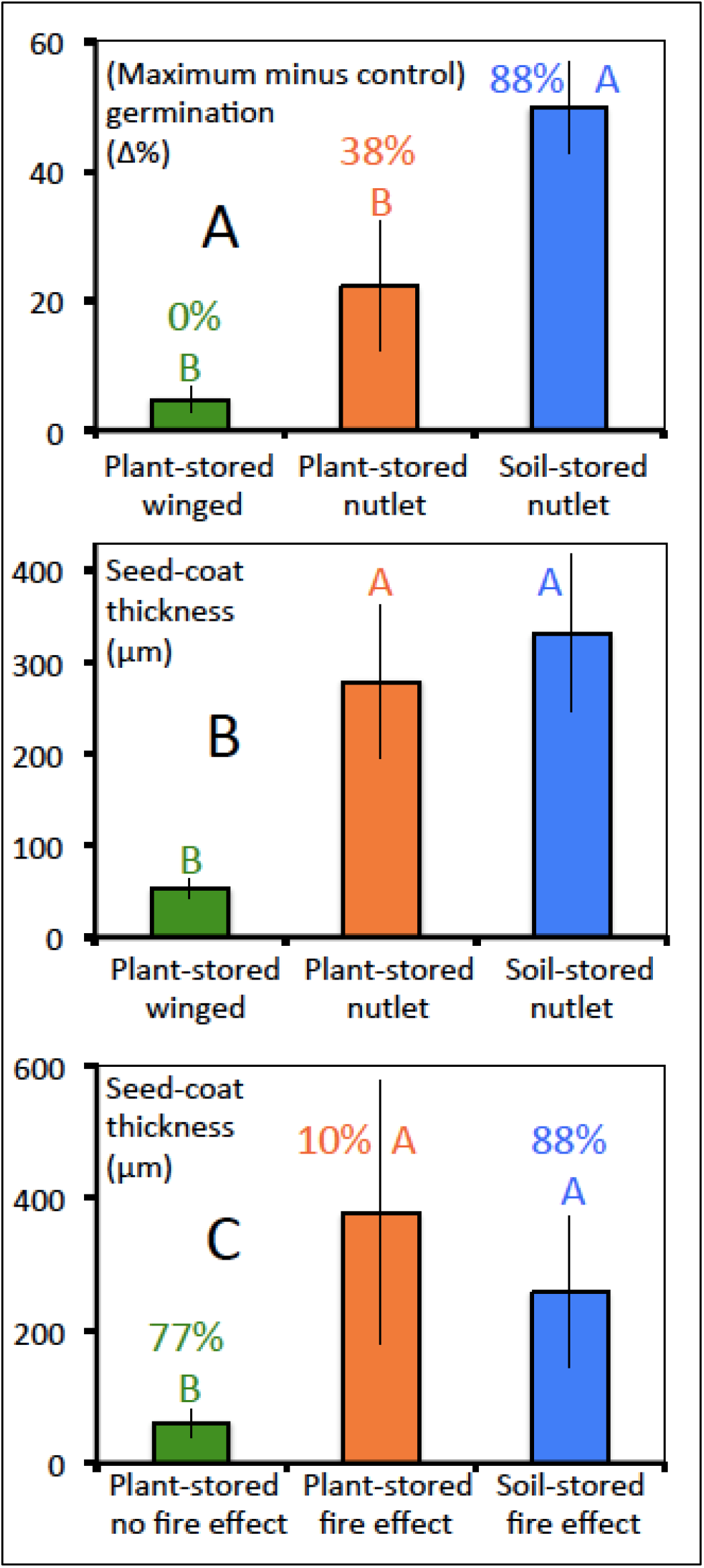
Means ± 95% CIs for all species in each of three seed categories based on storage location and fruit morphology, plus results for ANOVA of weighted means followed by Tukey’s HSD test – letters significantly different at *P* < 0.01. A. (Maximum value among the three firetype treatments minus control) germination levels plus % of species showing biologically significant differences within each category; B. Seed-coat thicknesses for the three categories; and C. Seed-coat thicknesses plotted against plant- or soil-stored seeds that do or do not have biologically significant germination levels in response to a fire (smoke) property plus % of species showing biologically significant differences within each category.

## Discussion

Among fire-prone seed plants generally, five fire-response dormancy-release/germination syndromes can be recognized (Table 3). *Leucadendron* has representatives in three of them: syndrome 1 with almost all species having plant-stored, winged, single-seeded fruits lacking any need for a fire property to stimulate germination, and these possessed thin seed coats; syndrome 2 with plant/soil-stored, nutlet-bearing species responding to smoke but not heat, with relatively thick seed coats; and syndrome 4 with a few soil-stored nutlet-bearing species responding non-additively to both smoke and heat, also with relatively thick seed coats. Overall, the thicker the seed coat, a) the lower the level of germination in the absence of fire-related properties, and b) the greater the germination response to fire-related properties, such that the difference between the controls and fire-treated seeds increased linearly with increase in seed-coat thickness (Results, Fig. 1). Smoke was far more effective at increasing germination levels among the species with thicker seed coats, such that the co-presence of heat made a negligible difference to the outcomes (Fig. 1b,c). Even so, fire-type heat (80°C for 20 min) alone increased the germination response to a minor extent (Fig. 1a).

Thus, seed-coat thickness is a reasonable predictor of the extent to which fire-related properties, especially smoke, will bring germination levels up to those of species that do not require heat or smoke (~100%). Water permeability is an inverse function of seed-coat thickness (as also shown by Noodén *et al*., 1985; Frączek *et al*., 2005), with winged achenes (seed-coat thickness 25-140 μm) much more permeable than nutlets (150-600 μm), such that seed permeability declined in a power-function manner (best-fit curve). Since the amount of smoke chemicals absorbed is a function of water uptake (Baxter *et al*., 1994), and thicker seed coats are less permeable (shown here), it raises the possibility that (at least some of) these promotive chemicals serve to increase permeability to water and/or oxygen (Brown and van Staden, 1973; Ghebrehiwot *et al*., 2008; Jain *et al*., 2008; Brits and Manning, 2019).

Of the ten species with biologically significant smoke responses, two (*L. elimense, L. thymifolium*) responded significantly to both smoke *and* heat. Heat and smoke can be distributed patchily during a fire (Auld and Bradstock, 1996) and the ability to respond to more than one germination property has been suggested to maximise the capability of seeds to sense the passage of a fire (Kenny, 2000; Morris, 2000). Once heated, the hard seeds of many legumes become permeable (Burrows *et al*., 2018) and will now germinate as soon as the soil is moist and cool, so that smoke sensitivity would be redundant. Smoke-sensitive seeds are stored in a permeable state that allows smoke chemicals after fire to enter them, so that now heat-sensitivity is redundant. However, it is possible for smoke chemicals to drift into unburnt or scorched patches through diffusion and leaching (Ghebrehiwot *et al*., 2013), where the seeds are unlikely to have received a heat treatment. However, patches already occupied by plants are unlikely to lead to recruitment because they are outcompeted by the plants already present (Lamont *et al*., 2019). Thus, a dual response is most likely to be adaptive when a) seeds are in patches that will receive both heat and smoke, and b) the response is additive, especially if it is synergistic (syndrome 5A/B in Table 2).

**Table 2.**
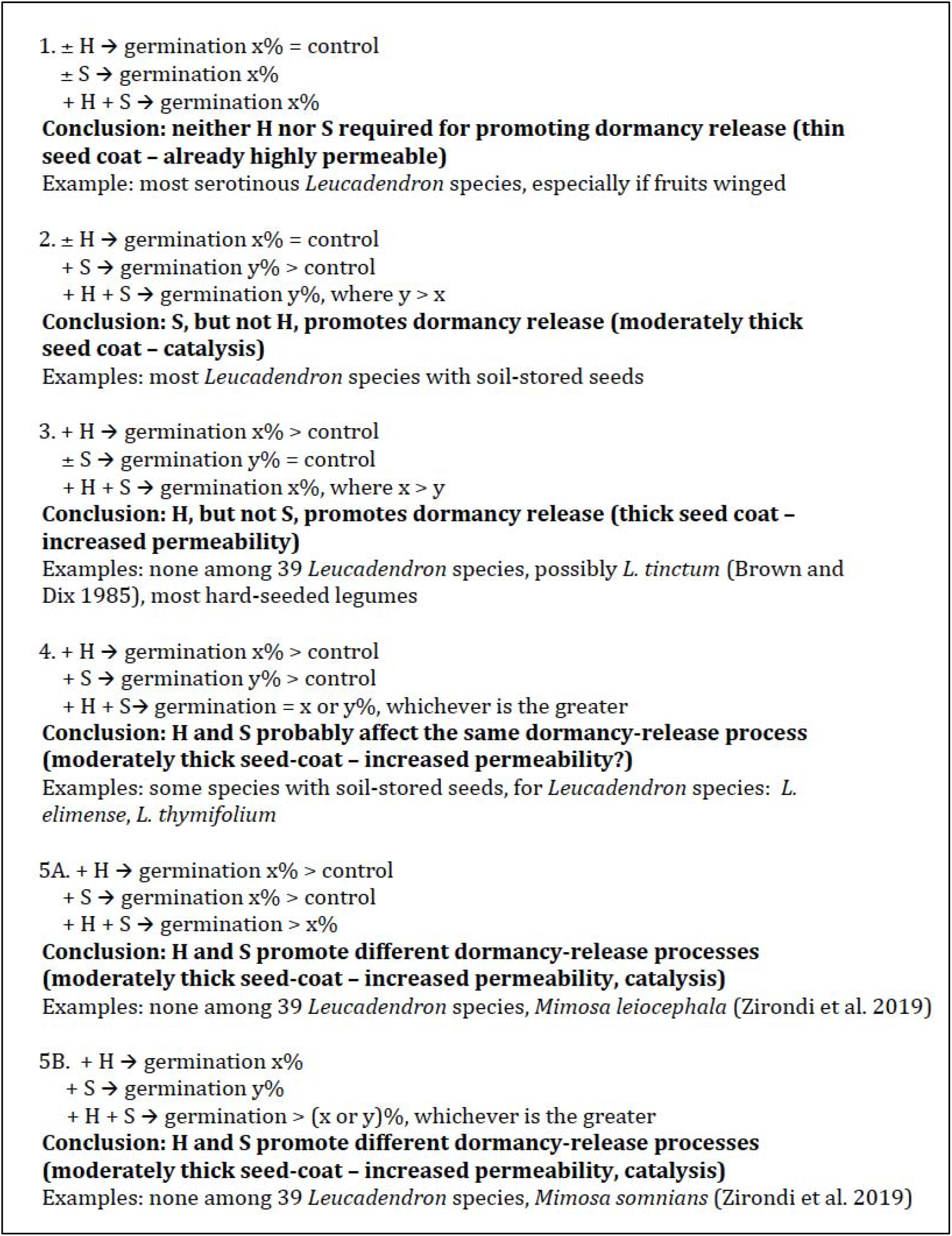
Five possible dormancy-release/germination syndromes based on individual and the additive effects of heat (H) and smoke (S) on germination that should apply to all fire-prone species, with examples from this study of 39 *Leucadendron* species (excluding *L. tinctum*) and elsewhere if not observed here. ± = presence or absence of fire component, → = leads to treatment outcome. () refers to likely mechanism of dormancy release as supported by relevant literature.

In a worldwide survey of 589 species subjected experimentally to both heat and smoke, 14.5% responded positively to both heat *and* smoke (BBL, unpublished) so that this phenomenon is not common but it is also not rare. In some cases, hard seeds become smoke-sensitive after they are heated and there is an additive or synergistic effect (Zirondi *et al*., 2019; syndrome 5A/B). The logical interpretation is that these environmental properties affect different processes that are well-known: heat renders the seeds permeable and smoke chemicals have a catalytic effect on the seed’s physiology. However, for *Leucadendron*, the two heat-responsive species were not impermeable to water, and their seed-coat thickness and permeability were not greater than some other smoke-responsive-only species (Fig. 2). Further, germination was no greater than with smoke *plus* heat than with smoke alone. But germination was greater than with heat alone that in turn was greater than for the controls.

These responses imply that smoke and heat may affect the same process (syndrome 4). Since heat probably increases permeability (no other function for a fire-type heat pulse on dormant seeds is known), and the thicker the seed coat the proportionately greater the smoke response, it seems in this case that heat supplements the permeability-enhancing role of smoke chemicals (Jain *et al*., 2008; Ghebrehiwot *et al*., 2008). Although difficult to envisage a relevant scenario, it appears that a non-additive response to smoke and heat is only likely to be adaptive when seeds receive fire-type heat in the absence of smoke. The improbability of such a situation might explain why this dual response is not better represented among fire-prone floras. This topic clearly needs further investigation, especially the relative role of heat and smoke in raising the permeability of non-hard seeds.

Our results also raise the interesting issue of *L. tinctum* that has >50% greater seed-coat thickness than the next thickest but germinated poorly despite high viability and did not respond to any treatment. In another study with this species, germination was raised from 12% to 45% with smoke and to 70% with smoke plus scarification (Brown and Botha, 2004), equivalent to syndrome 5 (Table 2). Brown and Dix (1985) showed that the seed coat was essentially a mechanical barrier to breaking dormancy by increasing germination from 20% to 80% after scarifying the seeds then covering them in lanolin. If our batch had exceptionally thick seed coats they could act like conventional ‘hard’ seeds and only respond to heat. It is possible then that our heat treatment (80°C for 20 min) was not ‘severe’ enough to scarify most seeds in this species, although the complete lack of a smoke response is not easily explained.

That it is not just an issue of winged achenes versus nutlets is demonstrated when the eight species with plant-stored nutlets are considered. These are as thick as the soil-stored nutlets on average but their response to smoke is much less (Fig. 3A, B). Two were as thin as the winged seeds (controls 99% germination) and six were on average 50 μm thicker than the mean of the soil-stored nutlets (controls 61% germination). The three plant-stored species showing a biologically significant response to smoke had seed coats five times thicker than the plant-stored species that did not benefit from smoke (Fig. 3C). So, the key to their germination requirements is seed-coat thickness rather than storage location or morphology. Thus, some plant-stored nutlets may be just as permeable as the winged achenes as they have similar seed-coat thicknesses (Fig. 2). This may have functional significance - the confines of the cone are much less hazardous for survival than in the soil, and germination can proceed readily following postfire release as with the winged seeds. However, the six species with thick-walled, plant-stored nutlets increase their options to remain soil-stored if they arrive in a microsite conducive to germination (Lamont et al. 2021) or, if unsuitable, to remain dormant until the fire.

Thick, weakly permeable seed coats serve to increase longevity and heat tolerance in the soil but then the seed must rely on a special fire-related property, smoke, as distinct from just cool wet winters, to signal (and respond to) the onset of ideal recruitment conditions. Thus, this genus possesses ideal taxa to pursue the mechanisms by which stimulatory smoke chemicals serve to break dormancy, as closely related species that do not require smoke also exist. This includes the possibility of serving to improve permeability (Ghebrehiwot *et al*., 2008; Jain *et al*., 2008) that needs further investigation. The role of fire-type heat is usually considered to be breaking the impermeability of hard seeds. Yet here we have instances of seeds with moderate permeability that can also benefit from heat that suggests it may also serve to further increase the permeability of seeds that are already (weakly) permeable. Evidence available so far indicates that this dual response to heat and smoke may exist among many hundreds of fire-prone species. This genus is ideal for pursuing the mechanisms by which seeds may respond to both smoke and heat and their functional significance.

## Acknowledgements

We thank a colleague who reviewed a draft of Newton *et al*. (2021) for suggesting the possible significance of seed-coat thickness in explaining our initial results, Berin Mackenzie for introducing the concept of biological significance to this work on which this follow-up research was based, and Richard Cowling and Tianhua He for early support.

## Author contributions

BBL and RJN conceived the project and designed the experiments; PGB performed all experiments and tests; BBL and PGB analysed the data; BBL and RJN wrote the draft manuscript; PGB contributed to drafts and all gave final approval for publication.

## Data availability

Data used for this analysis are given in Table 1 here and Table S5 of Newton *et al*. (2021).

